# CCR4 blockade leads to clinical activity and prolongs survival in a canine model of advanced prostate cancer

**DOI:** 10.1101/2021.04.12.439476

**Authors:** Shingo Maeda, Tomoki Motegi, Aki Iio, Kenjiro Kaji, Yuko Goto-Koshino, Shotaro Eto, Namiko Ikeda, Takayuki Nakagawa, Ryohei Nishimura, Tomohiro Yonezawa, Yasuyuki Momoi

## Abstract

Targeting regulatory T cell (Treg) infiltration is an emerging strategy for cancer immunotherapy. However, the efficacy of this strategy in advanced prostate cancer remains unclear. Here, we describe the therapeutic efficacy of this strategy in a canine model of advanced prostate cancer. We used dogs with naturally occurring prostate cancer to study the molecular mechanism underlying Treg infiltration into tumor tissues and the effect of anti-Treg treatment. We found that tumor-infiltrating Tregs were associated with poor prognosis in dogs bearing spontaneous prostate cancer. RNA sequencing and protein analyses showed that Treg infiltration was mediated by interaction between the tumor-producing chemokine, CCL17, and the receptor CCR4 expressed on Tregs. Dogs with advanced prostate cancer responded to mogamulizumab, a monoclonal antibody targeting CCR4, with improved survival and low incidence of clinically relevant adverse events. Exploratory analyses showed urinary CCL17 concentration and BRAF^V595E^ mutation to be independently predictive of the response to mogamulizumab. Analysis of a publicly available transcriptomic dataset of human prostate cancer showed that the CCL17/CCR4 axis correlated with the Treg marker, Foxp3. In silico survival analyses showed that high expression of CCL17 was associated with poor prognosis. Immunohistochemistry confirmed that tumor-infiltrating Tregs expressed CCR4 in human patients with prostate cancer. These findings suggest that anti-Treg treatment through the blocking of CCR4 is a promising therapeutic approach for advanced prostate cancer.

**One Sentence Summary:** Targeting regulatory T cell infiltration by CCR4 blockade induces objective responses and improves survival in a canine model of prostate cancer.

## Introduction

Prostate cancer is the most common malignancy in men, with an estimated 1.2 million new cancer cases and 359,000 deaths annually worldwide (*1*). Androgen deprivation therapy is a first-line treatment for advanced prostate cancer. This treatment alone, or in combination with chemotherapy, is initially effective in approximately 80%–90% of advanced prostate cancer cases. However, the disease eventually progresses to castration-resistant prostate cancer (CRPC) within months or years (*2*). Metastatic CRPC (mCRPC) has no curative treatment options and is associated with a poor prognosis. Although several treatments have been approved for mCRPC after progression with docetaxel chemotherapy, new treatment options that provide durable disease control are still needed.

Foxp3-expressing regulatory T cells (Tregs) play a role not only in the suppression of immune response against self-antigens but also in tumor progression by inhibiting the antitumor immunity. In humans, Treg infiltration has been observed in certain tumor tissues and has been associated with the progression of cancer and prognosis (*3, 4*). High infiltration of Tregs into tumor tissues correlates with poor prognosis in patients with melanoma, hepatocellular carcinoma, lung carcinoma, ovarian cancer, breast cancer, pancreatic cancer, and prostate cancer (*3, 5, 6*). In tumor-bearing mice, depletion of Tregs by administration of anti-CD25 enhances the antitumor immunity and leads to tumor eradication (*7, 8*). Anti-Treg therapy is under investigation for human patients with melanoma, adult T cell leukemia/lymphoma, lung cancer, and esophageal cancer (*9–11*). However, the role of Tregs and the therapeutic efficacy of its depletion in prostate cancer remain unclear.

Although rodent models are indispensable in cancer research, the controlled environment under which highly inbred rodents are kept offers completely different settings from the diverse conditions prevalent in human cancer. Mice-based preclinical studies have often failed to predict the outcomes of human clinical trials (*12*). The average rate of successful translation from rodent models to clinical cancer trials is less than 10% (*13*). In fact, clinical trials in patients with mCRPC using cytotoxic T lymphocyte antigen 4 (CTLA-4) or programmed cell death protein 1 (PD-1)/PD-L1 inhibitors have been less satisfactory, with limited survival benefit when administered as a monotherapy in unselected patients, than mice-based preclinical studies (*14–16*). Mouse models of prostate cancer are inadequate because of the lack of several features, such as genetic heterogeneity, molecular complexity, immune responses, symptoms, disease progression, metastatic behavior, and long-term evaluation. More relevant animal models of advanced prostate cancer are warranted to study the disease.

The canine prostate gland shares morphological and functional similarities with the human prostate. Dogs are the only animals to present with a significant incidence of spontaneous prostate cancer, with clinical features, including late age at onset and metastatic patterns resembling those in humans (*17, 18*). Therefore, naturally occurring prostate cancer in companion dogs could serve as a bridge between laboratory animal models and human patients. Unlike humans, most dogs with prostate cancer present with an advanced and aggressive disease. Local invasion to the urethra, bladder trigone, and ureter is common and causes obstruction of urine outflow leading to hydronephrosis. Metastases have been reported in >40% of dogs at diagnosis and in approximately 80% at death, with spread primarily to locoregional lymph nodes, lungs, liver, and bone (*19*). Dogs with prostate cancer often develop bone metastases in the lumbar vertebra, pelvis, and/or femur, associated with pain and neurological deficits (*19*). An important difference between human and canine prostate cancer is the role of androgen. Canine prostate cancer usually does not respond to androgen deprivation therapy or surgical castration (*20*). Interestingly, the cancer occurs more frequently in castrated male dogs (*18*). Dog-based clinical trials can be conducted in a comparatively shorter duration because dogs have a shorter life span than humans. Thus, the canine prostate cancer model could be informative to study the pathogenesis of advanced prostate cancer, especially mCRPC, and to assess biomarkers and therapeutics.

Here, we show that tumor-infiltrating Tregs are associated with poor prognosis in dogs with spontaneous prostate cancer, and that Treg migration is mediated by C-C chemokine ligand 17 (CCL17) and the receptor, CCR4. We also demonstrate the therapeutic potential of anti-CCR4 treatment in dogs with prostate cancer. Furthermore, we show that the CCL17/CCR4 axis is associated with Treg infiltration and poor prognosis in human prostate cancer patients. The canine model of advanced prostate cancer paves the way for the translation of the CCR4 blockade therapy to human patients with prostate cancer.

## Results

### Tumor-infiltrating Tregs associate with poor prognosis in canine prostate cancer

In certain tumors of dogs as well as of humans, Treg infiltration is associated with poor prognosis. We evaluated the abundance of tumor-infiltrating Tregs and the association between their density and prognosis in dogs with naturally occurring prostate cancer. Tissue samples were obtained from 18 dogs with prostate cancer (table S1); all these dogs underwent radical cystoprostatectomy. According to the World Health Organization (WHO) TNM classification for canine prostate cancer (*21*), 1/18 (6%) tumors were classified as T1 (intracapsular tumor, surrounded by normal gland), 2/18 (11%) as T2 (diffuse intracapsular tumor), 8/18 (44%) as T3 (tumor extending beyond the capsule), and 7/18 (39%) as T4 (tumor fixed, or invading neighboring structures). Nodal metastasis was detected in seven (39%) dogs. No distant metastasis (to the lungs or bones) was observed. After radical cystoprostatectomy, five dogs received no treatment, 10 received non-steroidal anti-inflammatory drugs (NSAIDs: piroxicam or carprofen), and two were administered chemotherapy (carboplatin or cyclophosphamide) in combination with NSAIDs.

We visualized the expression of Foxp3 using immunohistochemistry and examined the localization and number of tumor-infiltrating Tregs. Only a few Foxp3^+^ Tregs were detected in the normal canine prostate, whereas they were observed in both intratumoral area and peritumoral stroma of canine prostate cancer (Fig. 1A). Compared with normal tissues, Tregs were more frequently detected in dogs with prostate cancer (Fig. 1B). Gene expression of the immunosuppressive cytokine, IL-10, was elevated in canine prostate cancer compared with that in normal controls (Fig. 1C).

**Fig. 1.**
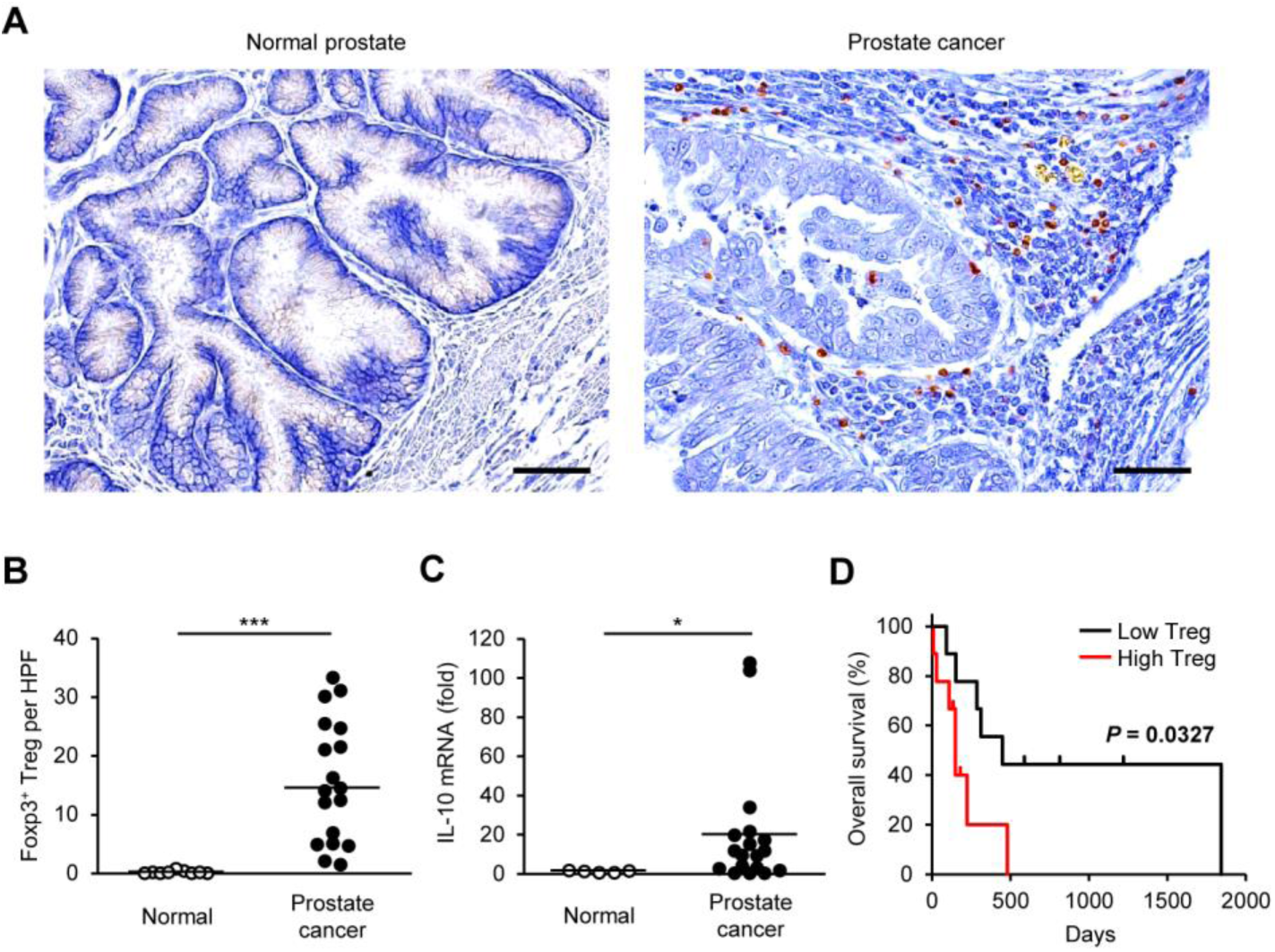
The number of tumor-infiltrating Tregs is associated with poor prognosis in dogs with prostate cancer. (**A**) Representative images of immunohistochemistry for Foxp3 in canine normal prostate and prostate cancer. Scale bar, 50 μm. (**B**) The number of Foxp3^+^ Tregs in the prostate of normal dogs (*n* = 9) and dogs with prostate cancer (*n* = 18). Median values are depicted by horizontal lines. \*\*\**P* < 0.001, nonparametric Mann– Whitney *U* test. (**C**) Expression of IL-10 mRNA in the prostate of normal dogs (*n* = 5) and dogs with prostate cancer (*n* = 18). Mean values are depicted by horizontal lines. \**P* < 0.05, nonparametric Mann–Whitney *U* test. (**D**) Kaplan–Meier curves of overall survival according to the number of intratumoral Tregs in dogs with prostate cancer (*n* = 18). Cases were classified as having a high or low density of Foxp3^+^ Tregs according to the median number. Log-rank test.

During the follow-up period, 13 of 18 dogs died (12 from progression of prostate cancer; 1 from disseminated intravascular coagulation). At the end of the study period, 5 dogs were alive. The median overall survival (OS) in dogs with prostate cancer was 201.5 days (range, 8–1841). Based on the median number, we classified each prostate cancer case as having a high or low density of Foxp3^+^ Tregs. The OS for cases with high Tregs was shorter than that for cases with low Tregs (Fig. 1D). These findings suggest that tumor-infiltrating Tregs alter the clinical outcome for dogs with prostate cancer.

### CCL17 expression is elevated in canine prostate cancer

To identify molecules inducing Treg infiltration in canine prostate cancer, we explored differentially expressed genes (DEGs) of chemoattractants for Tregs by RNA-Seq analysis. The analysis revealed several genes that were differentially regulated, with a *P*-value < 0.01, in canine prostate cancer when compared with normal controls. In total, 4,599 DEGs showed significant changes between normal and prostate cancer tissues. Of these, 2,301 DEGs were upregulated and 2,298 were downregulated in canine prostate cancer (Fig. 2A). Sixteen chemokine genes were upregulated in canine prostate cancer compared with normal prostate tissues (Fig. 2B). Quantitative RT-PCR showed approximately 700-fold increase in the expression of CCL17 gene in prostate cancer compared with the expression in normal canine prostate (Fig. 2C). Urinary CCL17 concentration was increased in dogs with prostate cancer (Fig. 2D). No significant difference was observed in serum CCL17 concentration between normal dogs and dogs with prostate cancer (Fig. 2E).

**Fig. 2.**
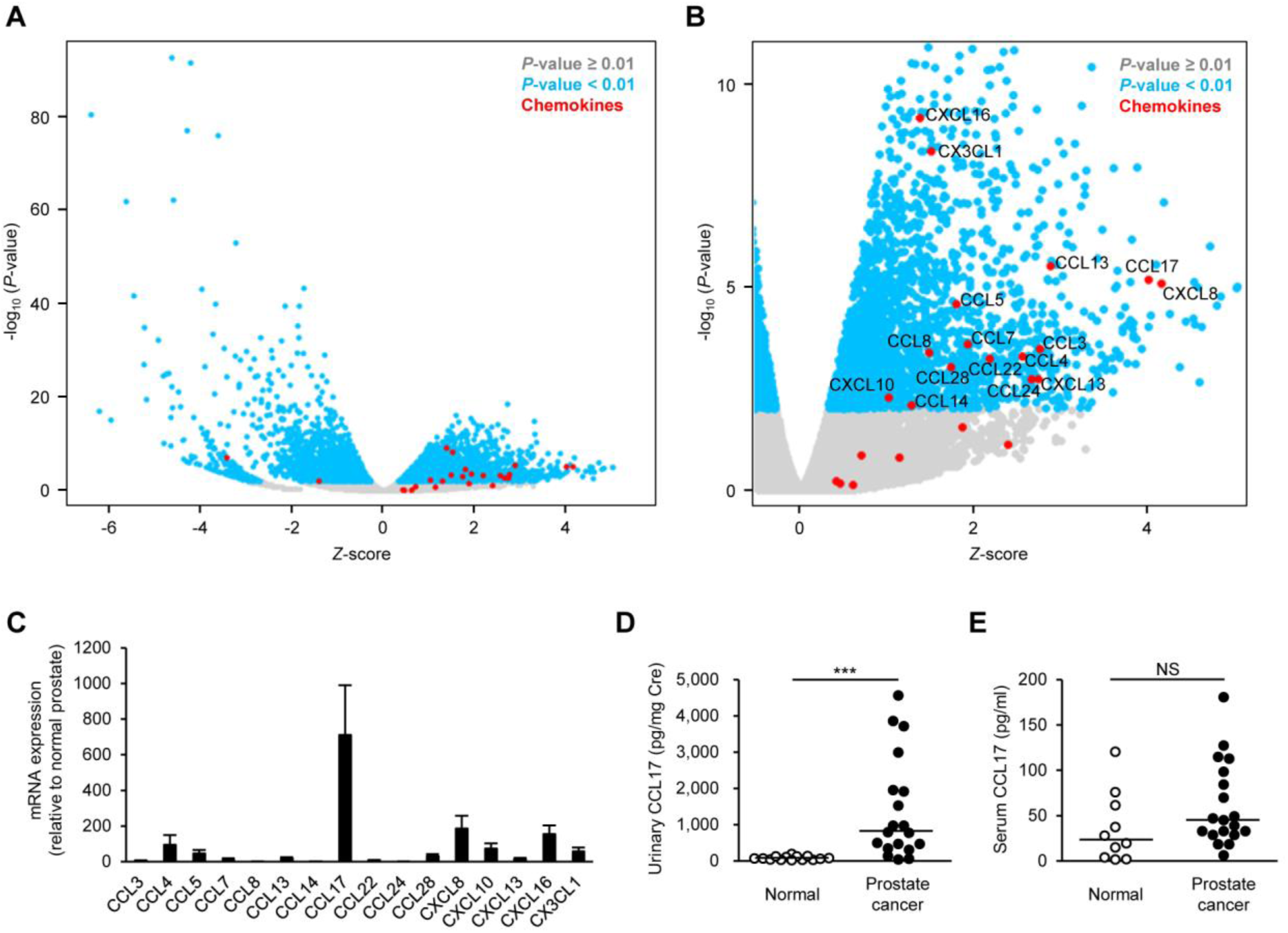
CCL17 expression is elevated in dogs with prostate cancer. (**A**) Volcano plot showing *Z*-score for differentially expressed genes between canine normal prostate (*n* = 4) and prostate cancer (*n* = 14) as determined by RNA-Seq. (**B**) Volcano plot showing *Z*-score for upregulated genes. Chemokine genes are enriched in upregulated genes. (**C**) Chemokine mRNA expression in canine prostate cancer (*n* = 18) relative to that in the normal prostate (*n* = 5) as determined by quantitative RT-PCR. (**D**) Urinary CCL17 concentration in normal dogs (*n* = 14) and in dogs with prostate cancer (*n* = 19). Mean values are depicted by horizontal lines. \*\*\**P* < 0.001, nonparametric Mann–Whitney *U* test. (**E**) Serum CCL17 concentration in normal dogs (*n* = 10) and in dogs with prostate cancer (*n* = 19). NS, not significant, nonparametric Mann–Whitney *U* test.

### Tumor-infiltrating Tregs express CCR4 in canine prostate cancer

CCL17 induces chemotaxis via the chemokine receptor, CCR4 (*22*). We examined whether CCR4-expressing cells infiltrate canine prostate cancer tissues using immunohistochemistry. As expected, CCR4^+^ cells with a mononuclear lymphoid morphology were abundant in prostate cancer but not in normal tissues (Fig. 3A). More CCR4^+^ cells were evident in dogs with prostate cancer (Fig. 3B), their density being positively correlated with the number of tumor-infiltrating Foxp3^+^ Tregs (Fig. 3C). Double immunofluorescence analysis confirmed that tumor-infiltrating Foxp3^+^ Tregs expressed CCR4 (Fig. 3D). Furthermore, CCR4^+^ Foxp3^-^ non-Treg cells were also present in the tumor tissues; however, CCR4^-^ Foxp3^+^ Tregs were rarely observed (Fig. 3D). The median proportions of CCR4^+^ Foxp3^+^ cells, CCR4^+^ Foxp3^-^ cells, and CCR4^-^ Foxp3^+^ cells were 38.1% (range, 19.4–50.2), 60.5% (49.5–79.4), and 0.7% (0–1.4), respectively, in dogs with prostate cancer. These observations suggest that infiltration of Tregs into the tumor tissue in canine prostate cancer is through the CCL17/CCR4 axis.

**Fig. 3.**
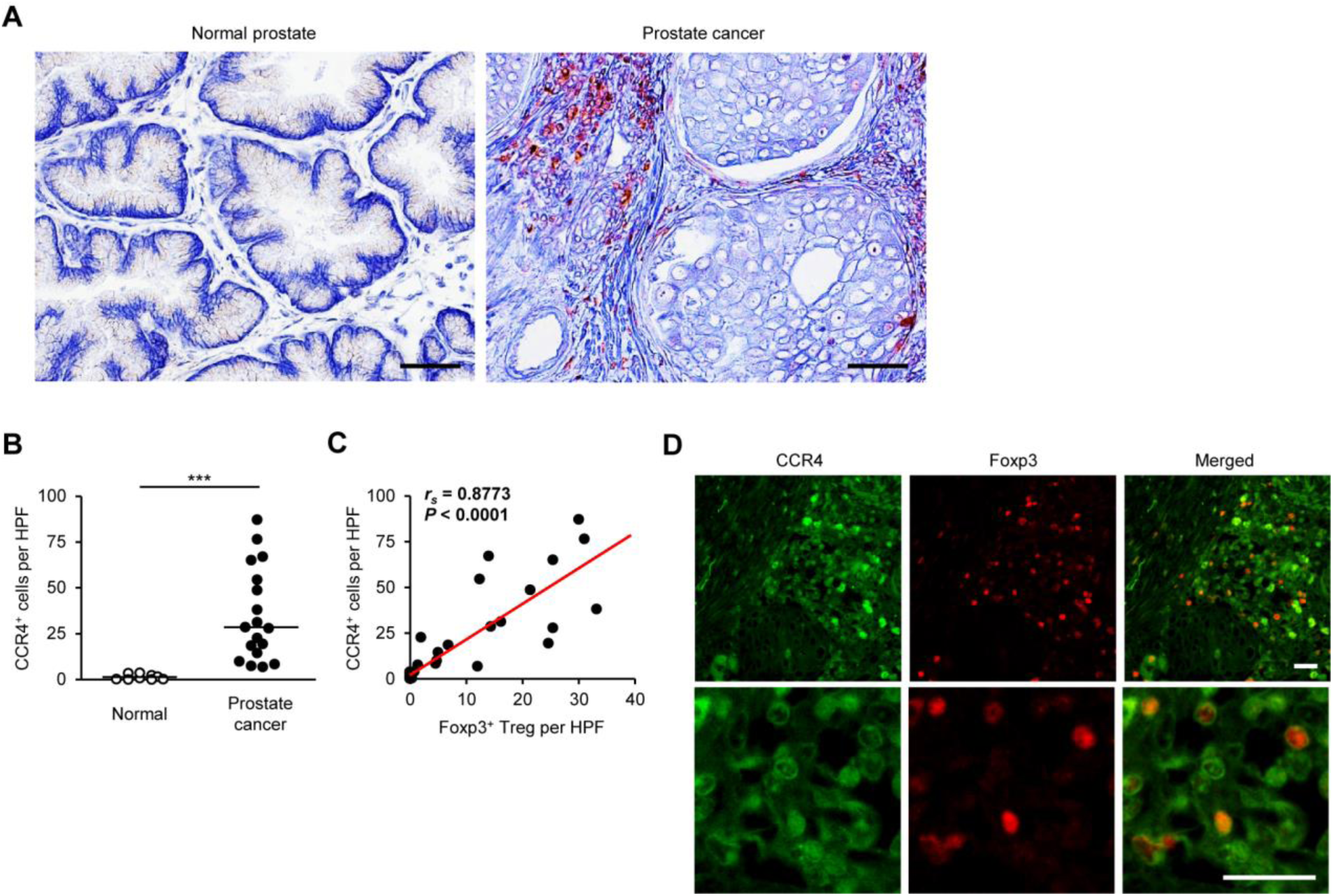
Tumor-infiltrating Tregs express CCR4 in dogs with prostate cancer. (**A**) Representative images of immunohistochemistry for CCR4 in canine normal prostate and prostate cancer. Scale bar, 50 μm. (**B**) The number of CCR4^+^ cells in the prostate of normal dogs (*n* = 9) and dogs with prostate cancer (*n* = 18). Median values are depicted by horizontal lines. \*\*\**P* < 0.001, nonparametric Mann–Whitney *U* test. (**C**) Correlation between Foxp3^+^ Tregs and CCR4^+^ cells in dogs with prostate cancer. Spearman rank correlation coefficient. (**D**) Representative images of immunofluorescence for CCR4 (green) and Foxp3 (red) in canine prostate cancer. Scale bar, 25 μm.

### BRAF^V595E^ mutation correlates with tumor-infiltrating Tregs and CCL17/CCR4 expression in canine prostate cancer

A somatic point mutation in the BRAF gene (BRAF^V595E^), which is homologous to the human BRAF^V600E^ mutation, is present in over 70% of dogs with bladder and prostate cancers (*23*). We have recently shown that BRAF^V595E^ mutation induces CCL17 production and contributes to Treg recruitment in dogs with bladder cancer (*24*). Thus, we investigated whether BRAF^V595E^ mutation influences tumor-infiltrating Tregs or the CCL17/CCR4 axis in dogs with prostate cancer. Of the 28 dogs with prostate cancer used in this study, BRAF^V595E^ mutation was detected in 21 (75%) cases (table S1). Tumor-infiltrating Foxp3^+^ Tregs and CCR4^+^ cells were increased in cases with BRAF^V595E^ mutation in comparison to that in cases with wild-type BRAF (fig. S1). Moreover, urinary CCL17 concentration in cases with the BRAF^V595E^ mutation was higher than in cases with wild-type BRAF. The BRAF^V595E^ mutation was not detected in any normal dog (table S1). Analysis of publicly available datasets of human prostate cancer showed that BRAF gene alterations (mutation, fusion, or copy number alteration) were found in 4.7%–6.5% and 3%–4% of patients with metastatic and nonmetastatic prostate cancer, respectively (fig. S2).

### Anti-CCR4 treatment leads to clinical responses and improves survival in spontaneous canine prostate cancer

A humanized anti-human CCR4, mogamulizumab, is commercially available for the treatment of CCR4^+^ adult T-cell leukemia/lymphoma (*9*). We previously confirmed that mogamulizumab crossreacts with canine CCR4 and depletes Tregs in dogs (*25*). To assess the clinical activity of the anti-CCR4 treatment in spontaneous canine prostate cancer, we compared 23 dogs that received mogamulizumab (1 mg/kg every 3 week) and piroxicam (0.3 mg/kg every 24 h) with a cohort of 23 age-, sex-, and tumor stage-matched dogs that received piroxicam alone (table S2). Typically, dogs that received mogamulizumab treatment in combination with piroxicam had a reduction in the tumor burden (Fig. 4A, upper and movie S1). Three dogs (case ID. M3, M13, and M15) exhibited fluid retention in the prostate due to necrosis of the tumor, although there was no reduction in the mass (Fig. 4A, lower and movie S2). Compared to dogs treated with piroxicam alone, those administered mogamulizumab/piroxicam treatment had a greater reduction in the size of the primary tumor (Fig. 4B). The median percentage of maximum tumor reduction in dogs treated with mogamulizumab/piroxicam and in those treated with piroxicam alone was –22.7% (range, –43.6% to 8.5%) and 2.9% (range, –34.7% to 95.9%), respectively. In 23 dogs with mogamulizumab/piroxicam, 7 (30%) obtained partial response (PR), 14 (61%) had stable disease (SD), and 2 (9%) had progressive disease (PD). In 23 dogs with piroxicam alone, 2 (9%) obtained PR, 13 (56%) had SD, and 8 (35%) had PD. The clinical response to mogamulizumab/piroxicam was higher than the response to piroxicam alone (Fig. 4C). At the cutoff time for the study data (March 1, 2021), 3 (13%) dogs that received mogamulizumab were alive. The median progression-free survival (PFS) in dogs treated with mogamulizumab/piroxicam and in those administered piroxicam alone was 189 (range, 21–573) days and 57 (range, 6–210) days, respectively. The median OS in dogs treated with mogamulizumab/piroxicam and in those treated with piroxicam alone was 296 (range, 86–1,000) days and 99 (range, 6–468) days, respectively. The PFS and OS in dogs treated with mogamulizumab in combination with piroxicam were longer than in those treated with piroxicam alone (Fig. 4D).

**Fig. 4.**
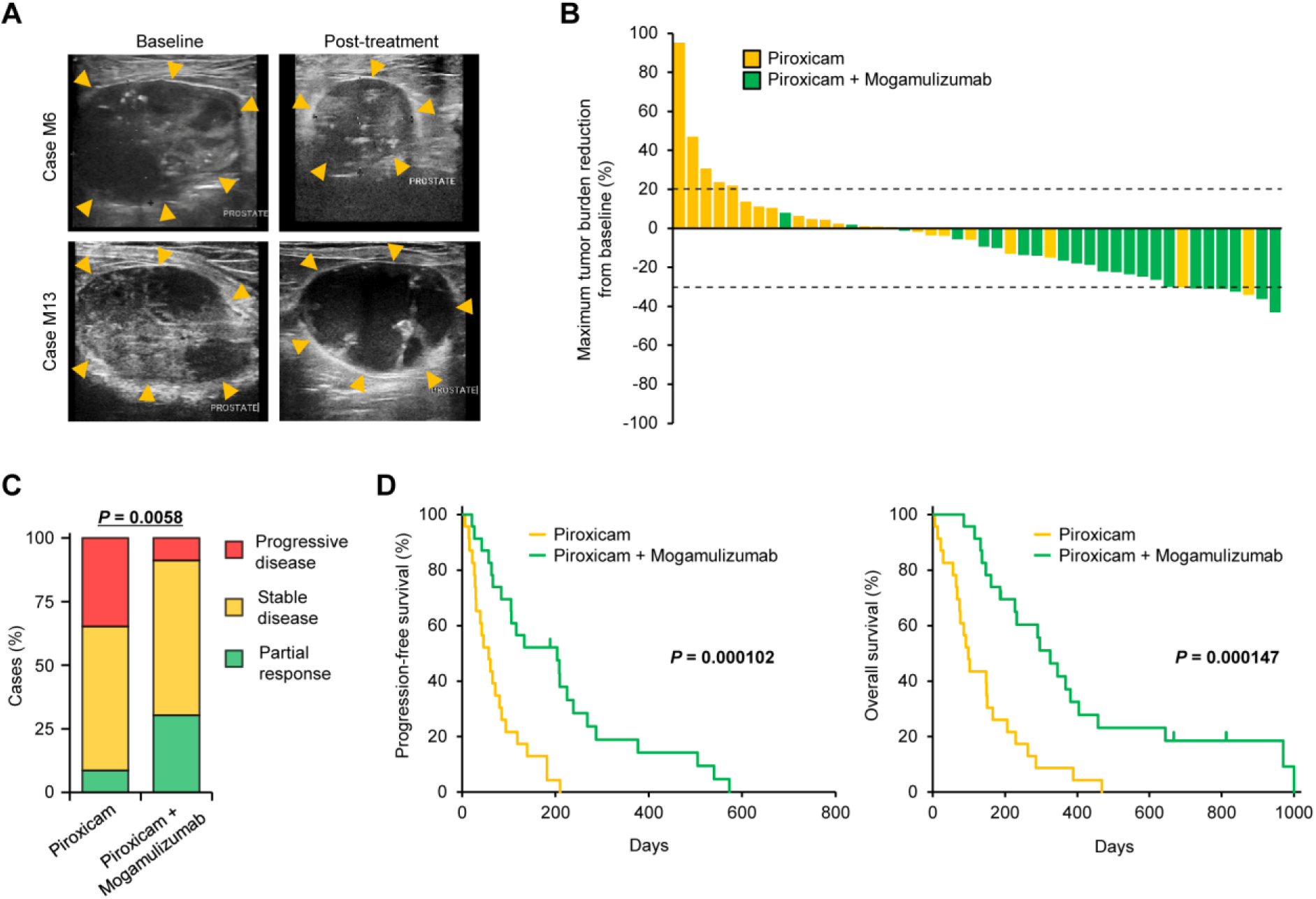
Anti-CCR4 therapy induces clinical responses and improves survival in dogs with prostate cancer. (**A**) Representative ultrasonographic images of prostate masses (arrowheads) in dogs treated with mogamulizumab and piroxicam. In case M6, the prostate mass shrunk after four cycles of treatment compared to baseline. In case M13, the prostate size did not change but necrosis was observed inside the mass after four cycles of treatment. (**B**) Waterfall plot showing the maximum percentage of tumor burden reduction from baseline in dogs treated with piroxicam (*n* = 23, yellow) or in dogs treated with mogamulizumab and piroxicam (*n* = 23, green). Dashed lines indicate –30% (partial response) and +20% (progressive disease). (**C**) Clinical responses in dogs treated with piroxicam (*n* = 23) or mogamulizumab and piroxicam (*n* = 23). Cochran–Armitage test. (**D**) Progression-free survival (left) and overall survival (right) in dogs treated with piroxicam (*n* = 23, yellow) or mogamulizumab and piroxicam (*n* = 23, green). Log-rank test.

Among 23 dogs treated with mogamulizumab/piroxicam, 17 (74%) had an adverse event (Table 1). All treatment-related adverse events were grade 1 or 2, and many were transient. The most frequent adverse events were increased alkaline phosphatase (35%), increased alanine transaminase (17%), vomiting (17%), and anorexia (13%). As suspected immune-related adverse events, pancreatitis (grade 2) and infusion reaction (grade 1) were observed in one case each (4%). Because mogamulizumab is a humanized antibody, the risk of allergic reactions in dogs was assumed; however, the allergic adverse events observed in this study were urticaria and rash only in one case (4%). No serious allergic reactions, such as anaphylaxis, were observed. There were no grade 3–5 treatment-related adverse events, and no dog exhibited an event leading to treatment withdrawal. Lymphopenia was observed in some cases (17% with grade 1 and 13% with grade 2), which was considered as the pharmacologic effect of mogamulizumab.

**Table 1.** Adverse events in dogs treated with mogamulizumab (n = 23)

| Event | Number of cases (%) |  |  |  |
| --- | --- | --- | --- | --- |
|  | Grade 1 | Grade 2 | Grade 3-5 | Total |
| Any event | 11 (47.8) | 9 (39.1) | 0 | 17 (73.9) |
| Nonhematologic |  |  |  |  |
| Increased ALP | 4 (17.4) | 4 (17.4) | 0 | 8 (34.8) |
| Increased ALT | 2 (8.7) | 2 (8.7) | 0 | 4 (17.4) |
| Vomiting | 2 (8.7) | 2 (8.7) | 0 | 4 (17.4) |
| Anorexia | 1 (4.3) | 2 (8.7) | 0 | 3 (13.0) |
| Lethargy/fatigue | 2 (8.7) | 0 | 0 | 2 (8.7) |
| Pancreatitis | 0 | 1 (4.3) | 0 | 1 (4.3) |
| Urticaria | 0 | 1 (4.3) | 0 | 1 (4.3) |
| Rash | 1 (4.3) | 0 | 0 | 1 (4.3) |
| Diarrhea | 1 (4.3) | 0 | 0 | 1 (4.3) |
| Infusion reaction | 1 (4.3) | 0 | 0 | 1 (4.3) |
| Hematologic |  |  |  |  |
| Lymphopenia | 4 (17.4) | 3 (13.0) | 0 | 7 (30.4) |

Taken together, these results suggest that the anti-CCR4 treatment leads to clinical responses and improves survival without severe adverse events in dogs with spontaneous prostate cancer.

### Urinary CCL17 and BRAF^V595E^ mutation are biomarkers for responses to the anti-CCR4 treatment

In a previous canine clinical trial of mogamulizumab for bladder cancer, urinary CCL17 was shown to be associated with clinical response (*25*). We investigated the association of pretreatment urinary CCL17 with the response in dogs with prostate cancer. No association was noted between urinary CCL17 and clinical response in dogs treated with piroxicam alone (Fig. 5A). In the cohort of mogamulizumab/piroxicam treatment, dogs with PR had more urinary CCL17 than did dogs with SD or PD (Fig. 5A). In dogs treated with piroxicam alone, PFS and OS for cases with high urinary CCL17 were shorter than those for cases with low urinary CCL17 (Fig. 5B). In contrast, high urinary CCL17 was associated with a longer OS in dogs treated with mogamulizumab/piroxicam (Fig. 5C).

**Fig. 5.**
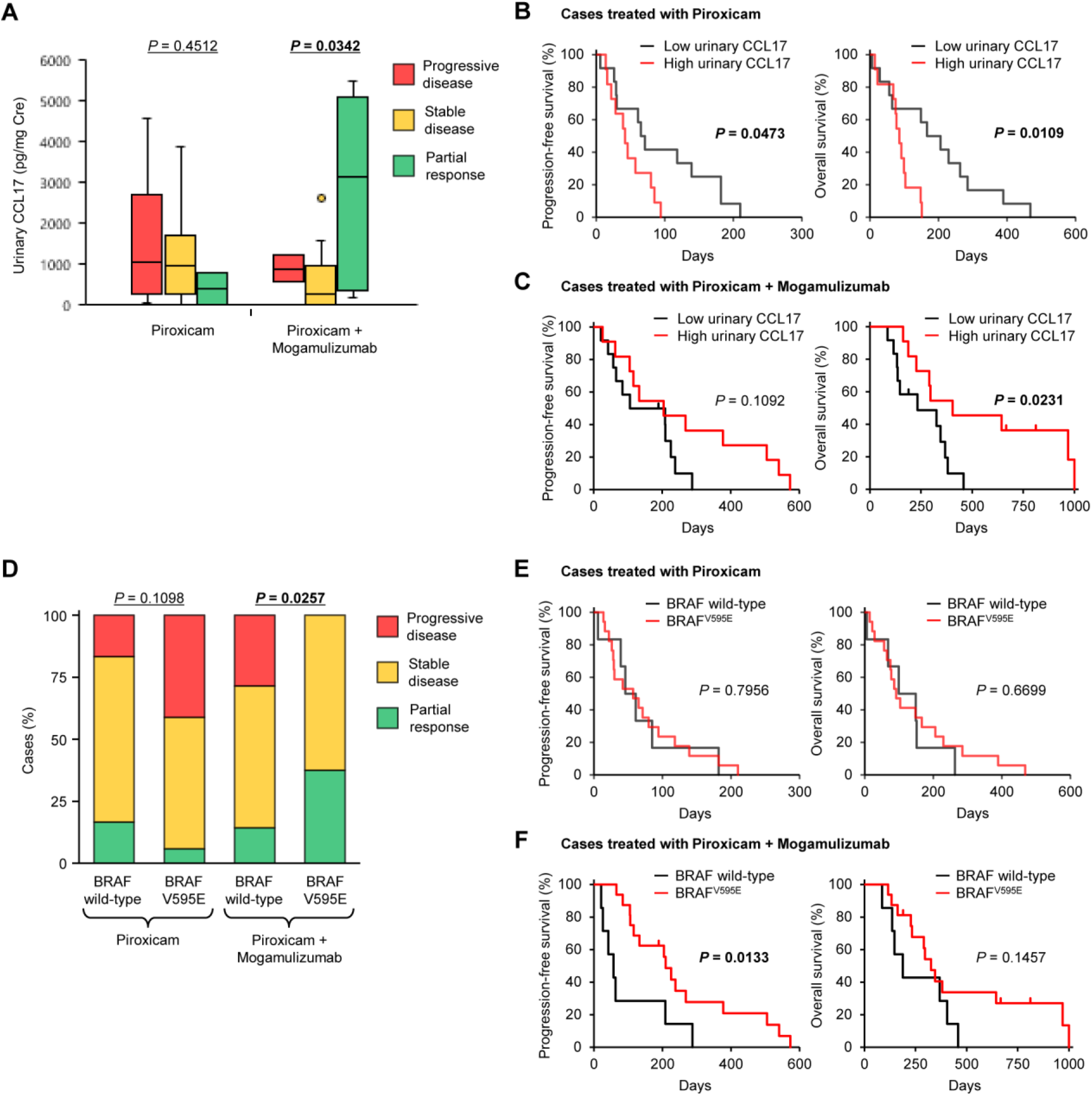
Urinary CCL17 concentrations and BRAF^V595E^ mutation are associated with clinical responses and outcomes of anti-CCR4 treatment. (**A**) Association of urinary CCL17 concentrations with response to treatment. Kruskal–Wallis test. (**B** and **C**) Kaplan–Meier curves of progression-free survival (left) and overall survival (right) according to urinary CCL17 concentrations in dogs treated with piroxicam (*n* = 23, **B**) or mogamulizumab and piroxicam (*n* = 23, **C**). Cases were classified as having a high or low concentration of urinary CCL17 according to the median. Log-rank test. (**D**) Association of BRAF^V595E^ mutation with response to treatment. Cochran–Armitage test. (**E** and **F**) Kaplan–Meier curves of progression-free survival (left) and overall survival (right) according to BRAF gene status in dogs treated with piroxicam (*n* = 23, **E**) or mogamulizumab and piroxicam (*n* = 23, **F**). Log-rank test.

We further examined the association between BRAF^V595E^ mutation and response. There was no association between BRAF^V595E^ mutation and clinical response in dogs treated with piroxicam alone, whereas BRAF^V595E^ mutation was associated with favorable response in dogs treated with mogamulizumab/piroxicam (Fig. 5D). Similarly, PFS and OS in dogs treated with piroxicam alone were not related to BRAF^V595E^ mutation (Fig. 5E). In dogs treated with mogamulizumab/piroxicam, PFS for cases with BRAF^V595E^ mutation was longer than that for cases with wild-type BRAF (Fig. 5F). These findings suggest that urinary CCL17 and BRAF^V595E^ mutation are useful biomarkers for predicting the clinical response and outcome to mogamulizumab treatment in dogs with prostate cancer.

### Human and canine prostate cancers exhibit common gene expression signatures

We hypothesized that transcriptional patterns of human and canine prostate cancers would be conserved. To systematically assess transcriptional patterns across species, we performed RNA-Seq analysis of canine prostate cancer and compared the data to a TCGA dataset of human prostate cancer (*26, 27*). We selected statistically significant DEGs (q < 0.01) between canine prostate cancer and normal tissues and extracted concordant genes in the expression data of humans. We identified 2,297 genes. The expression patterns of these genes in canine and human prostate tissues were visualized with t-SNE (fig. S3). Gene expression patterns in normal canine and human prostate samples were clearly distinct, whereas prostate cancer samples were not clearly divided by species. These results suggest that similarities in gene expression signatures in a subset of prostate cancer might share biology across species.

### CCL17/CCR4 axis associates with tumor-infiltrating Tregs and poor prognosis in human prostate cancer

Given the promising outcomes and favorable clinical responses of the anti-CCR4 treatment in the comparative canine trial, we examined whether the CCL17/CCR4 axis is associated with tumor-infiltrating Tregs and prognosis in human prostate cancer patients. We searched for mRNA expression of CCL17, CCL22, CCR4, and Foxp3 in the publicly available transcriptomic dataset of human prostate cancer from TCGA PanCancer Atlas (*n* = 493). High mRNA expression of CCL17, CCL22, CCR4, and Foxp3 was detected in 5%, 3%, 2.2%, and 5% of patients with prostate cancer, respectively (Fig. 6A). We found a correlation between mRNA expression of Foxp3 and CCL17 (*r*_s_ = 0.44, *P* = 9.0 × 10^-25^), CCL22 (*r*_s_ = 0.58, *P* = 3.9 × 10^-46^), and CCR4 (*r*_s_ = 0.60, *P* = 4.7 × 10^-50^; Fig. 6B). Immunohistochemistry in serial sections of human tissues revealed mononuclear lymphoid cells stained positive for Foxp3 and CCR4 in prostate cancer but not in normal prostate (Fig. 6C). Compared to the normal prostate, Foxp3^+^ Tregs and CCR4^+^ cells were more frequent in patients with prostate cancer (Fig. 6D). The density of Foxp3^+^ Tregs was positively correlated with CCR4^+^ cells (Fig. 6E). Double-labeling immunofluorescence confirmed that Foxp3^+^ tumor-infiltrating Tregs expressed CCR4 (Fig. 6F). To evaluate whether the ligands of CCR4 are associated with prognosis in human prostate cancer, we performed in silico survival analyses using TCGA dataset. We identified that high CCL17, but not CCL22, expression is associated with shorter PFS (Fig. 6G). These findings suggest that anti-CCR4 may have therapeutic value for the treatment of human prostate cancer.

**Fig. 6.**
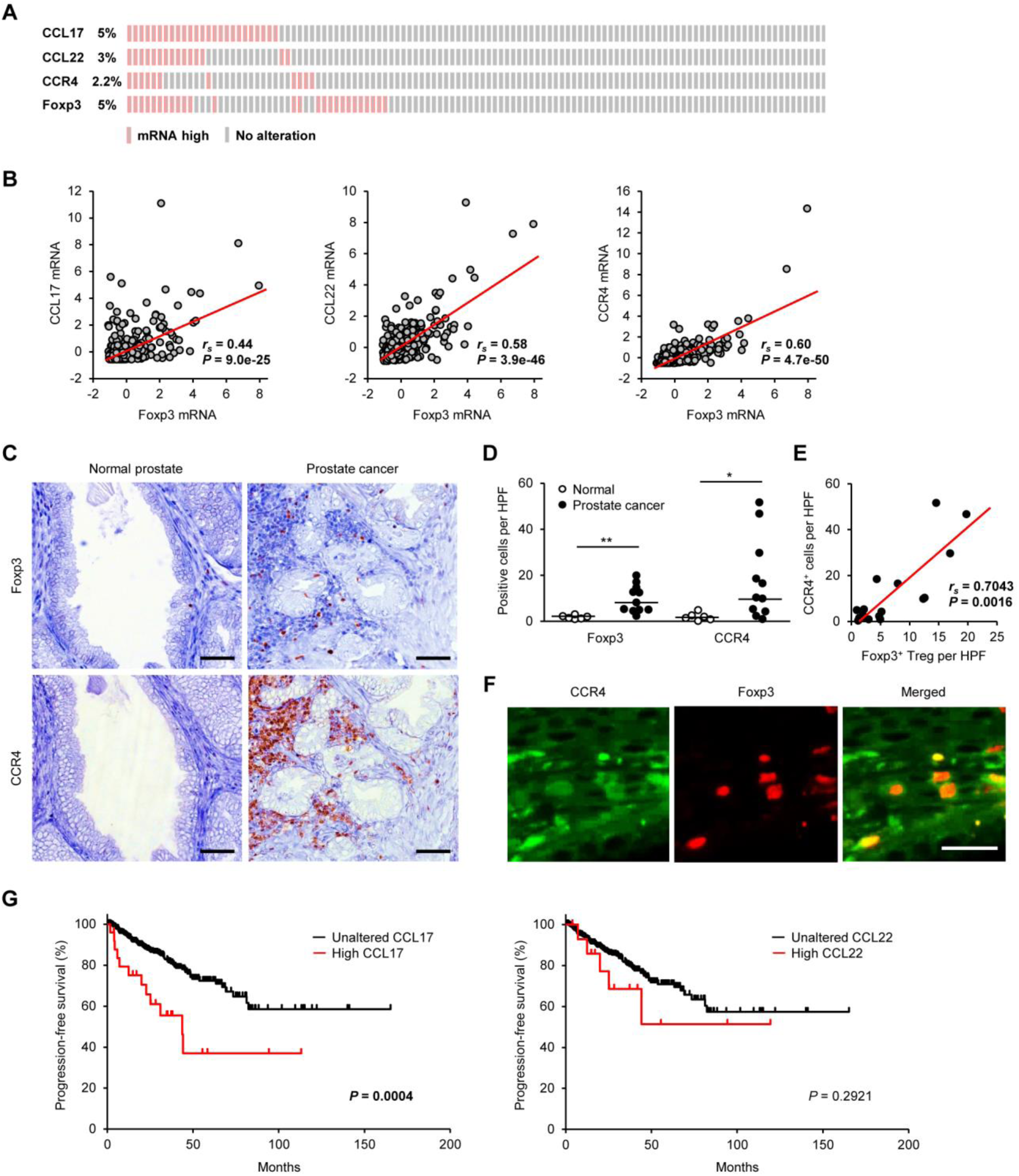
The CCL17/CCR4 axis is associated with tumor-infiltrating Tregs and prognosis in human patients with prostate cancer. (**A**) Expression data for CCL17, CCL22, CCR4, and Foxp3 mRNAs in human prostate cancer samples from TCGA dataset (*n* = 493). (**B**) Linear regression analysis between Foxp3 expression and CCR4-associated genes (CCL17, CCL22, and CCR4) in human prostate cancer samples from TCGA dataset (*n* = 493). Spearman rank correlation coefficient. (**C**) Representative images of immunohistochemistry for Foxp3 and CCR4 in human prostate cancer. Scale bar, 50 μm. (**D**) The number of Foxp3^+^ Tregs and CCR4^+^ cells in the prostate of normal volunteers (*n* = 6) and patients with prostate cancer (*n* = 11). Median values are depicted by horizontal lines. \**P* < 0.05, \*\**P* < 0.01, nonparametric Mann–Whitney *U* test. (**E**) Correlation between Foxp3^+^ Tregs and CCR4^+^ cells in patients with prostate cancer. Spearman rank correlation coefficient. (**F**) Representative images of immunofluorescence for CCR4 (green) and Foxp3 (red) in canine prostate cancer. Scale bar, 25 μm. (**G**) Kaplan–Meier curves of progression-free survival according to the mRNA expression of CCL17 (left) and CCL22 (right) in human patients with prostate cancer from TCGA dataset (*n* = 493). Log-rank test.

## Discussion

Immunotherapy is rapidly transforming cancer treatment across a range of tumor types. Prostate cancer tissues often contain immune cells, suggesting that this cancer is a target of host antitumor immunity (*28*). In 2010, sipuleucel-T, a cancer vaccine that targets prostatic acid phosphatase, was approved by FDA for the first immunotherapy of mCRPC (*29*). However, other immunotherapeutic approaches, including immune checkpoint inhibitors, have not been successful against mCRPC to date (*15, 16*). This failure may be explained by highly immunosuppressive microenvironment in prostate cancer (*30, 31*). An alternative approach is needed to evoke antitumor immunity and conquer the immunosuppressive microenvironment. In this study, we have shown that Foxp3^+^ Tregs infiltrate tumor tissues through the CCL17/CCR4 pathway, and CCR4 blockade exerts an antitumor effect in dogs with advanced prostate cancer. In our canine clinical trial, objective response rates (ORR) of mogamulizumab in combination with piroxicam treatment were 30% (7 of 23 dogs), and the median PFS and OS were 189 days and 296 days, respectively. Although there is limited literature regarding treatment of canine prostate cancer, these results compare well with tests of piroxicam alone in this study (ORR, 9%; median PFS, 57 days; median OS, 99 days) or chemotherapy (carboplatin, mitoxantrone, or cyclophosphamide) in combination with NSAIDs (ORR, 4%; median PFS, 89 days; median OS, 106 days) (*32*). The reported median OS in dogs with prostate cancer following prostatectomy ranges from 19 to 231 days (*33–35*), indicating a relatively higher therapeutic efficacy of the mogamulizumab/piroxicam treatment. We also found that CCL17/CCR4 pathway is associated with Treg infiltration and poor prognosis in human prostate cancer. These findings collectively suggest the potential of mogamulizumab for the treatment of human prostate cancer. Whether CCR4 blockade can enhance the clinical efficacy of sipuleucel-T and immune-checkpoint inhibitors warrants investigation.

The mogamulizumab treatment was well tolerated in dogs. Most treatment-related adverse events were of grade 1 or 2. Vomiting and anorexia were manageable with systemic antiemetic treatment alone. Allergic reactions, such as rash and urticaria, were observed but were mild, and only one case was treated with diphenhydramine. No intervention was required in other cases exhibiting allergic reactions. Consistent with the pharmacologic effect of mogamulizumab, 30% of dogs had a reduction in lymphocyte counts from pretreatment levels; however, there was no increased risk of clinically threatening infections. In humans, the most common adverse events associated with mogamulizumab treatment are infusion reactions (21%– 89%), skin rashes (51%–63%), chills (23.8%–59%), and nausea (19%–31%), which are manageable and reversible (*9, 36, 37*). These safety data, including a lack of renal toxicity, suggest that patients with advanced prostate cancer, who are often elderly and predisposed to renal impairment, may be better able to tolerate mogamulizumab treatment than chemotherapy.

We show that the likelihood of CCR4 blockade therapy response (tumor burden reduction and survival) can be increased by determining the urinary CCL17 concentration or BRAF^V595E^ somatic mutation. These findings indicate a link between the BRAF^V595E^ mutation and Treg recruitment via the CCL17/CCR4 pathway in dogs with prostate cancer. In humans, BRAF gene mutations occur in up to 8% of all cancers (*38*). Melanoma has the highest frequency of BRAF mutations (approximately 40%–60% of patients), with the majority harboring a BRAF^V600E^ mutation. Herein, we confirmed that BRAF gene alterations occur in up to 6.5% of human patients with prostate cancer. Given that more than 70% of canine prostate cancers harbor BRAF^V595E^ mutation, the companion dog model could be a highly relevant platform for comparative cancer research to understand molecular events leading to BRAF mutation and further develop a BRAF-targeted therapy. It will be interesting to investigate whether BRAF inhibitors can enhance the antitumor effect of the CCR4 blockade therapy.

Among in vivo models for cancer study, mouse models represent the most widely used system. The advantages of mouse models include the ease of genetic manipulation, short gestation period, and low maintenance cost. Although mouse models have remained valuable in the preclinical testing of anticancer agents, the major challenges in considering mouse models as a translational platform are the lack of replicating tumor heterogeneity, genetic diversity, and microenvironment, which are hallmarks of human cancers. The use of conditional systems, inducible systems, and patient-derived xenografts has partially offset this limitation; however, improvements are yet to be made to address the issues of tumor microenvironment and inter-patient variability in treatment response observed in the clinical setting (*39*). On the contrary, naturally occurring canine cancers better resemble human cancers with regard to heterogeneity, clinical sign, histopathology, disease progression, and metastatic behavior (*40, 41*). Comparative oncology clinical trials in companion dogs play a growing role in cancer research and drug development efforts. In particular, an intact immune system and natural co-evolution of tumor and microenvironment support exploration of novel immunotherapeutic strategies (*41*). Herein, we show that the CCL17/CCR4 axis associates with Treg infiltration into the tumor tissues in both canine and human prostate cancer. Other canine malignancies, including non-Hodgkin lymphoma, glioma, bladder cancer, breast cancer, lung cancer, melanoma, and osteosarcoma, share genotypic and phenotypic similarities with human counterparts (*41*). In 2017, the US National Cancer Institute (NCI) Cancer Moonshot initiative, named PRECINCT (Pre-medical Cancer Immunotherapy Network for Canine Trials), funded several canine clinical trials for accelerating the subsequent application of novel immunotherapeutic approaches to humans. Studies on canine spontaneous cancer provide an opportunity for comparative research and drug development linking basic studies with human clinical trials.

In conclusion, we demonstrate that the CCL17/CCR4 pathway associates Treg infiltration into tumor tissues and adverse outcomes in canine and human prostate cancer. CCR4 blockade leads to clinical activity and improves survival without severe toxicity profiles in a canine model of advanced prostate cancer. These findings suggest that anti-CCR4 treatment may be a viable strategy for reducing immunosuppression caused by Tregs, thereby augmenting antitumor immunity in both dogs and humans.

## Materials and Methods

### Study design

The overall objective of this study was to assess the therapeutic benefit of anti-Treg treatment in advanced prostate cancer using a naturally occurring canine model. A comparative study in companion dogs with spontaneous prostate cancer was used to bridge preclinical and human studies. The study was designed to (i) explore the molecular mechanism of Treg infiltration into the tumor tissues, (ii) examine whether the blockade of the identified pathway (CCL17/CCR4) would result in an efficient clinical response to spontaneous prostate cancer in a canine clinical trial, and (iii) validate the relationship of the CCL17/CCR4 pathway with Treg infiltration and survival in human patients with prostate cancer. The phase II canine clinical trial was conducted at the Veterinary Medical Center of the University of Tokyo (VMC-UT). The client-owned dogs with prostate cancer were treated with anti-CCR4 therapy in combination with piroxicam (the standard drug for canine prostate cancer). Age-, sex-, and tumor stage-matched dogs with prostate cancer treated with piroxicam alone were used as a control cohort for clinical response and survival. No placebo control, blinding, or randomization was performed in the study. The histological and bioinformatic analyses of human prostate cancer were performed using human tissues and publicly available datasets, respectively.

### Ethical statements

The study protocol of tumor tissue sampling from client-owned dogs and canine clinical trial were approved by the Animal Care and Clinical Research Committees of the VMC-UT (approval no. VMC2016-02). Written informed consent was obtained from all dog owners. All experimental methods were performed in accordance with the approved guidelines. All human tissue samples were collected per the U.S. Common Rule, and human clinical data were collected per the HIPAA guidelines.

### Canine prostate cancer model, selection criteria, and sample collection

Characteristics of dogs used for histological, gene expression, and protein expression analyses are presented in table S1. For histological analysis, archival formalin-fixed, paraffin-embedded prostate cancer tissues were obtained from 18 dogs at the VMC-UT. All dogs underwent radical cystoprostatectomy. The diagnosis of prostate cancer was confirmed by histopathology. Normal canine prostate tissues were obtained from nine healthy beagles euthanized for another experimental purpose. Survival time and current status (alive, deceased, or lost) of all dogs were determined by medical record or interview. OS was defined as the time from radical cystoprostatectomy to the established cause of death of the animal at the end of the study (April 9, 2019).

Fresh tumor tissues from 18 dogs with prostate cancer were analyzed for gene expression (table S1). Archival snap-frozen normal prostate tissues collected from five healthy beagles were used as control (table S1). For enzyme-linked immunosorbent assay (ELISA), serum samples were collected from 19 dogs with prostate cancer and 10 healthy dogs (table S1). Fresh urine samples were collected using a urethral catheter from 19 dogs with bladder cancer and 14 healthy dogs (table S1).

Dogs with spontaneous prostate cancer (*n* = 23) were enrolled in a canine clinical trial of anti-CCR4 treatment. As control dogs, 23 age-, sex-, and tumor stage-matched dogs with prostate cancer, treated with piroxicam during the study period, were recruited. Dogs that had been administered chemotherapy or radiation therapy were excluded from this clinical trial. Before treatment, tumor stage was defined as per the WHO criteria for canine prostate cancer (*21*). Characteristics of dogs used in the clinical trial are summarized in table S2.

### Human samples

We purchased paraffin-embedded sections of prostate specimens from 6 healthy volunteers and 11 patients with prostate cancer (OriGene Technologies). Characteristics of human patients with prostate cancer are summarized in table S3.

### Immunohistochemistry

The expression of Foxp3 and CCR4 was examined by immunohistochemistry with 4 μm-thick paraffin-embedded sections (*25*). Heat-induced antigen retrieval was performed by autoclaving the sections for 5 min at 121°C in 10 mM sodium citrate buffer (pH 6.0). Endogenous peroxidase activity was blocked by incubation with REAL Peroxidase-Blocking Solution (Dako) at room temperature for 10 min. The sections were blocked with 5% skim milk in Tris-buffered saline with 0.1% Tween 20 (TBST) at room temperature for 60 min and then incubated with primary antibodies, rat anti-Foxp3 (1:400 dilution, clone FJK-16s, eBioscience) or mouse anti-CCR4 (1:100 dilution, clone 1G1, BD Biosciences), at 4°C overnight. Secondary antibodies were applied as follows: EnVision polymer reagent for mouse (Dako) at room temperature for 45 min or a biotin-labeled anti-rat IgG (Vector Laboratories) at 37°C for 30 min followed by HRP-labeled streptavidin (Dako) at room temperature for 30 min. The reaction products were visualized with 3,3′-diaminobenzidine (DAB). For double immunofluorescence, antigen retrieval, endogenous peroxidase blocking, and milk blocking were performed as described above. Primary antibodies, rat anti-Foxp3 (1:100 dilution, clone FJK-16s, eBioscience) and mouse anti- CCR4 (1:100 dilution, clone 1G1, BD Biosciences), were applied at 4°C overnight. Immunofluorescence was performed using secondary antibodies, Alexa Fluor 594 goat anti-rat IgG (1:500, Invitrogen) and Alexa Fluor 488 donkey anti-mouse IgG (1:500, Invitrogen). Images were captured using a fluorescence microscope (BZ-X800; Keyence).

For human prostate tissues, antigen retrieval, endogenous peroxidase blocking, and milk blocking were performed as described above. As primary antibodies, rabbit anti-human Foxp3 (1:100 dilution, clone 1054C, R&D Systems) or mouse anti-human CCR4 (1:100 dilution, clone 1G1, BD Biosciences), was applied at 4°C overnight. Immunohistochemistry was performed by using secondary antibodies, EnVision polymer reagent for rabbit or mouse (Dako) at room temperature for 45 min, followed by DAB detection. Immunofluorescence was performed using secondary antibodies, Alexa Fluor 598 goat anti-rabbit IgG (1:500, Invitrogen) and Alexa Fluor 488 donkey anti-mouse IgG (1:500, Invitrogen).

Cells with clear lymphocyte morphology, distinct nuclear staining for Foxp3 or cytoplasmic staining for CCR4 were evaluated as positive. Foxp3^+^ or CCR4^+^ cells were quantified in 10 representative fields of each slide (40× magnification) using the ImageJ software (*42*).

### Canine prostate cancer RNA-Seq

Total RNA was extracted from prostate cancer and normal prostate tissues using the RNeasy Mini Kit (Qiagen). RNA integrity was examined with an Agilent 2100 Bioanalyzer (Agilent Technologies) and RNA integrity number (RIN) values of all samples were >7. Sequencing libraries were prepared with the TruSeq Stranded mRNA Library Prep Kit for NeoPrep (Illumina). RNA-Seq (75 bp paired-end) was conducted using NextSeq 500 (Illumina) with the High Output Kit (Illumina), and a minimum of 35 million read-pairs was generated for each sample.

Quality controls and adaptor trimmings of fastq files for each sample were performed using the Trim Galore software based on FastQC and Cutadapt (version 0.6.3; https://www.bioinformatics.babraham.ac.uk/projects/trim_galore/). Trimmed fastq data were mapped to canine genomes (CanFam3.1) by STAR (ver. 2.7.3a) (*43*) and transcript abundance was estimated using RSEM (ver 1.3.3) (*44*) with gene transfer file for Ensembl (CanFam3.1.98; https://www.ensembl.org). These gene count data were used to normalize and extract differential gene expressions with an EdgeR-based R package, TCC (ver. 1.26.0) (*45*). The results of normalized expression gene data between two groups were scaled to have a mean = 0 and SD = 1 (z-score) by R package, genefilter (ver. 1.68.0), and visualized using a volcano plot by R (ver. 3.6.1).

The datasets used and/or analyzed during the current study are available from the corresponding author on reasonable request and will also be available at the DDBJ Sequenced Read Archive repository (https://www.ddbj.nig.ac.jp/index-e.html) with accession number DRA011773.

### Bioinformatic analysis of human prostate cancer

Datasets for human metastatic or nonmetastatic prostate cancer were accessed and BRAF gene alterations were analyzed through cBioPortal (*26, 27, 46-52*). A normalized mRNA expression dataset for human prostate cancer (TCGA, PanCancer Atlas) was accessed and downloaded from the cBioPortal and used to evaluate associations between CCL17, CCL22, CCR4, and Foxp3 expression. This data set includes mRNA profiles for 493 patients with prostate cancer (*26*). PFS analysis was done for CCL17 and CCL22 transcripts for the cases. Detailed information of prostate cancer patients including pathology diagnosis, clinical stage, and survival data can be downloaded from the cBioPortal website (https://www.cbioportal.org/).

### Comparative genomic analysis between dogs and humans

RNA-seq count data for human prostate samples were obtained from GDC data portal (https://portal.gdc.cancer.gov/). Five hundred fifty count data for HTseq analyzed (498 prostate cancer and 52 normal prostate tissues) from TCGA-PRAD project were downloaded for this study (*53*). These data were also analyzed in the same way as canine data.

The most statistically significant DEGs between canine prostate cancer and normal tissues (q < 0.01) were extracted. These canine genes were converted to HGNC symbols using BioMart (*54*) and then the expression data of concordant genes between dogs and humans were extracted. This analysis resulted in 2,297 gene symbols, which were then used for t-distributed stochastic neighbor embedding (t-SNE) analysis (Rtsne package; ver. 0.15) of all samples (*55*).

### Quantitative RT-PCR

We quantified mRNA expression levels of IL-10 and chemokines identified in the RNA-Seq analysis using 2-step real-time RT-PCR (Thermal Cycler Dice Real Time System, Takara Bio). The ribosomal protein L13a (RPL13A) and RPL32 were used as reference genes. The primer pair sequences are shown in table S4.

### ELISA

We measured canine CCL17 levels in the supernatants of serum and urine samples using the Canine TARC/CCL17 ELISA kit (Cusabio), according to the manufacturer’s protocol. Urinary creatinine concentration was measured using the LabAssay Creatinine kit (Wako), and urinary CCL17 was expressed as pg/mg of creatinine (Cre).

### Canine clinical trial design and interventions

Anti-CCR4 mAb (mogamulizumab, 1 mg/kg; Kyowa Hakko Kirin) was administered to dogs with metastatic or nonmetastatic prostate cancer in 30 min intravenous infusions once every 3 week. The dosage and administration interval were based on a previous study (*25*). Treatment was continued until dogs experienced disease progression, had unacceptable toxicity, or their owners stopped adhering to the study protocol. Piroxicam (0.3 mg/kg; Pfizer), the standard drug used to treat canine prostate cancer, was administered every 24 h starting at the time of mogamulizumab treatment. We used age-, sex-, and tumor stage-matched dogs with prostate cancer, treated with only piroxicam, as matched controls.

### Clinical assessment

Dogs were evaluated for clinical responses and toxicity at least once every 3 week by owner observations, physical exam, complete blood counts, serum chemical profiles, 3-view thoracic and 2-view abdominal radiography, and abdominal ultrasonography. A single ultrasound operator measured prostate masses following a standardized mapping procedure (*56*). For each dog, longitudinal and transverse views of the prostate were obtained, and three measurements (height, width, and longitudinal length) were recorded. We used ultrasound for imaging because it could be conducted without general anesthesia (as would be required for computed tomography or magnetic resonance in dogs). According to the canine response evaluation criteria in solid tumors (*57*), we defined the tumor response as follows: complete remission (CR; no cancer detected), partial response (PR; ≥30% decrease in the sum of the longest diameters of target lesions from baseline and no new tumor lesions), progressive disease (PD; ≥20% increase in the sum of the longest diameters or the development of new tumor lesions), and stable disease (SD; not meeting the criteria for CR, PR, or PD). PFS was defined as the time from the start of treatment until PD or death at the end of the study (March 1, 2021), and OS as the time from the start of treatment until death of the animal at the end of the study. Adverse events were assessed and classified according to the Veterinary Cooperative Oncology Group (VCOG) criteria (*58*).

### Biomarkers

For biomarker assessment, we collected fresh urine samples before treatments for CCL17 measurements and BRAF^V595E^ mutation. Urinary CCL17 was assessed by ELISA, as described above. BRAF^V595E^ mutations were examined by digital PCR assay using genomic DNA isolated from urine sediments, as previously described (*24*).

### Statistical analysis

All data in bar graphs are presented as mean ± SEM. We used the JMP Pro version 15.0 (SAS Institute) for statistical analyses. The Mann–Whitney *U* test was used for comparison between two groups. The Kruskal–Wallis test, followed by Dunn test, was used for multiple comparisons. Correlation between two variables was evaluated using the Spearman rank correlation coefficient. The Cochran–Armitage test for trend was used to evaluate clinical response and treatment or BRAF^V595E^ mutation. Survival curves were generated using the Kaplan–Meier method and compared using the log-rank test. Statistical significance was defined as *P* < 0.05.

## Supplementary Materials

Fig. S1. BRAF^V595E^ mutation is associated with intratumoral Foxp3^+^ Tregs, CCR4^+^ cells, and urinary CCL17 concentration in dogs with prostate cancer.

Fig. S2. BRAF gene alterations in human prostate cancer.

Fig. S3. Cross-species t-SNE analysis of canine and human prostate tissues.

Table S1. Clinical features of dogs with prostate cancer and factors analyzed in this study.

Table S2. Characteristics of dogs with prostate cancer in the clinical trial.

Table S3. Characteristics of human patients with prostate cancer.

Table S4. Primer pair sequences used for quantitative RT-PCR.

Movie S1. Representative ultrasonographic images of a prostate mass in case M6 treated with mogamulizumab.

Movie S2. Representative ultrasonographic images of a prostate mass in case M13 treated with mogamulizumab.

## Acknowledgments

We thank Drs. M. Tsuboi, J.K. Chambers, and K. Uchida for the histopathological review of the canine samples. Computations were partially performed on the NIG supercomputer at ROIS National Institute of Genetics. We would like to acknowledge and thank the canine patients, the owners, and the clinical care team at the Veterinary Medical Center of the University of Tokyo.

## Author contributions

SM conceptualized and designed the study. SM, TM, AI, KK, YGK, SE, and NM conducted the experiments. SM and TM analyzed the data. SM performed statistical analyses. TN, RN, TY, and YM provided administrative, technical, and material support. TY and YM supervised the study. SM and TM wrote the draft manuscript. All authors approved the final manuscript.

## Data and materials availability

Raw sequence data for RNA-Seq of canine tumor-normal pairs have been deposited in the DDBJ Sequenced Read Archive repository, (https://www.ddbj.nig.ac.jp/index-e.html) with accession no. DRA011773.

