## Supplementary Materials for "CCR4 blockade leads to clinical activity and prolongs survival in a canine model of advanced prostate cancer"

**The PDF file includes:**

Fig. S1. $BRAF^{V595E}$ mutation is associated with intratumoral $Foxp3^+$ Tregs, $CCR4^+$ cells, and urinary CCL17 concentration in dogs with prostate cancer.

Fig. S2. BRAF gene alterations in human prostate cancer.

Fig. S3. Cross-species t-SNE analysis of canine and human prostate tissues.

Table S1. Clinical features of dogs with prostate cancer and factors analyzed in this study.

Table S2. Characteristics of dogs with prostate cancer in the clinical trial.

Table S3. Characteristics of human patients with prostate cancer.

Table S4. Primer pair sequences used for quantitative RT-PCR.

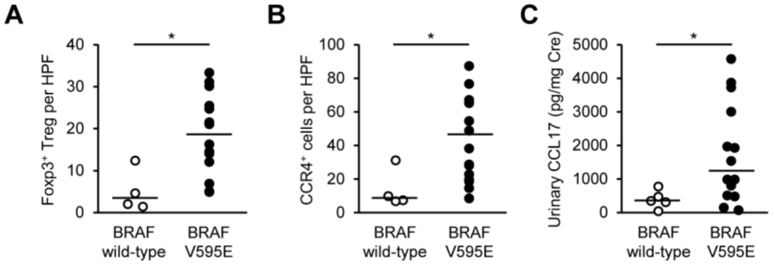


**Fig. S1. BRAF$^{V595E}$ mutation is associated with intratumoral Foxp3$^{+}$ Tregs, CCR4$^{+}$ cells, and urinary CCL17 concentration in dogs with prostate cancer.** (**A**–**C**) The number of intratumoral Foxp3$^{+}$ Tregs (**A**), CCR4$^{+}$ cells (**B**), and urinary CCL17 concentrations (**C**) in canine prostate cancer cases with wild-type BRAF and BRAF$^{V595E}$ mutation. $*P < 0.05$, nonparametric Mann–Whitney $U$ test.

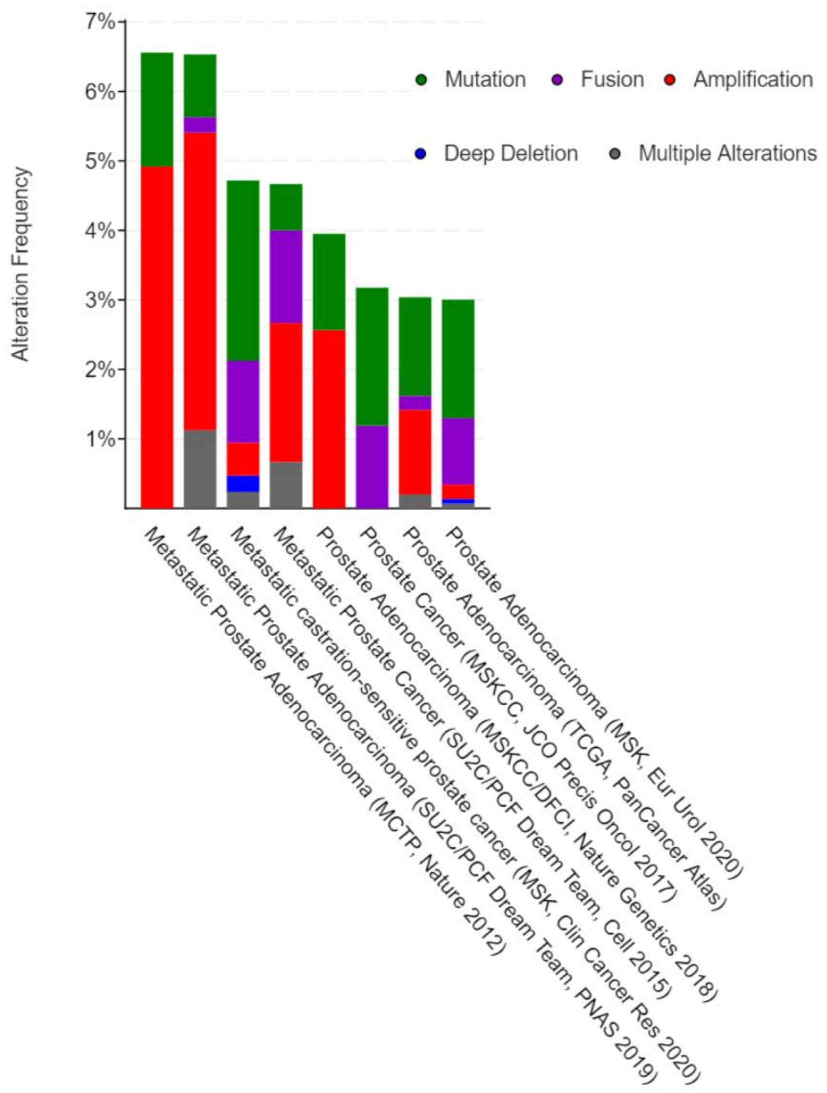


**Fig. S2. BRAF gene alterations in human prostate cancer.** Histogram displaying frequency of BRAF gene somatic mutation, fusion, copy-number amplification, deletion, and multiple alterations across eight human prostate cancer genomic datasets.

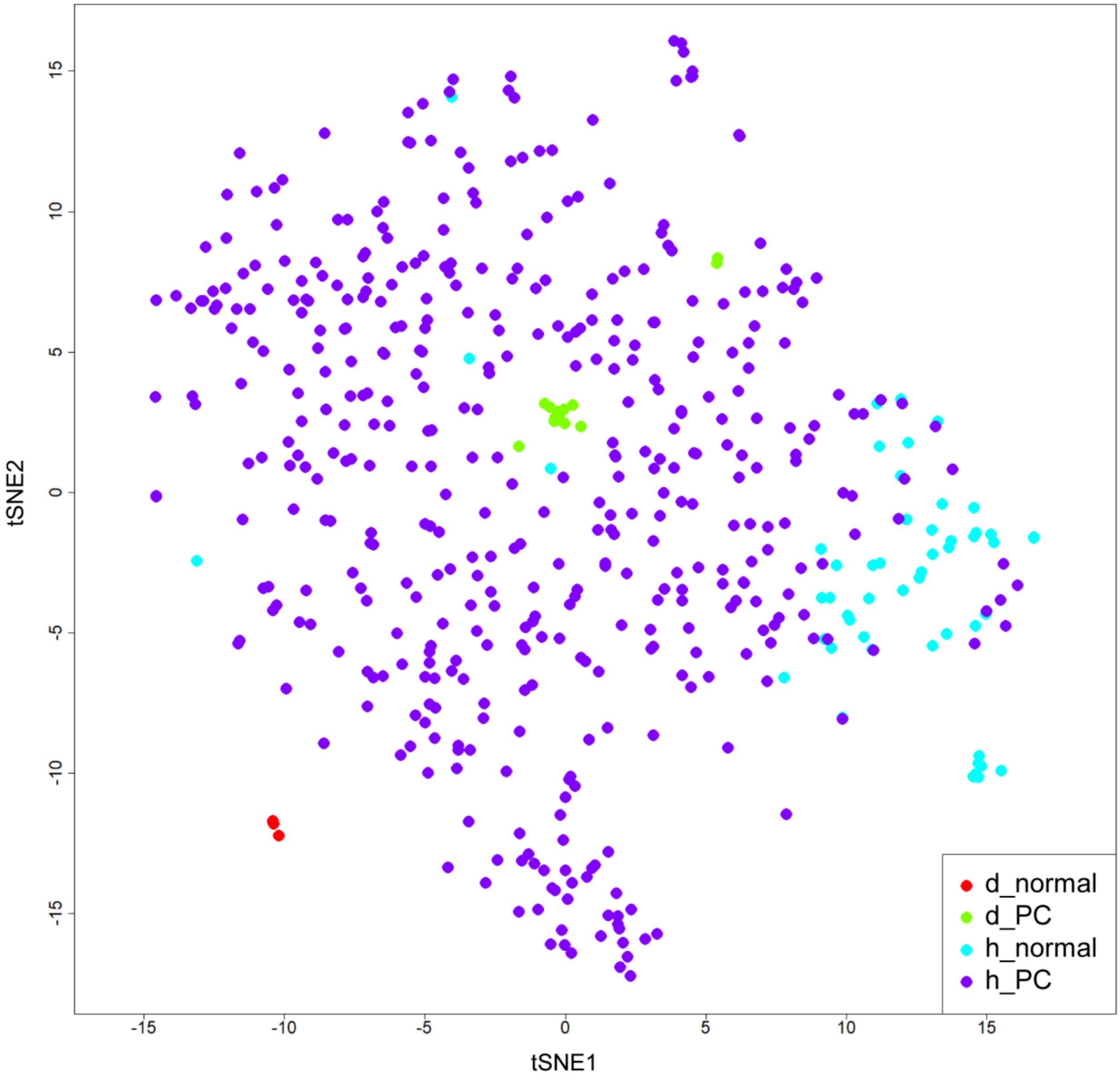


**Fig. S3. Cross-species t-SNE analysis of canine and human prostate tissues.** Differentially expressed genes (q < 0.01) between canine prostate cancer and normal prostate were selected and extracted concordant genes in the expression data of humans. The expression patterns of 2,297 differentially expressed genes in the cross-species tissues were visualized with t-SNE. Color-coded sphere showed each type of prostate tissue as following; red: canine normal prostate, green: canine prostate cancer, blue: human normal prostate, purple: human prostate cancer.

**Table S1. Clinical features of dogs with prostate cancer and factors analyzed in this study.**

| Case ID | Clinical features | | | | | | Analyzed factors[†] | | | | | |
|---|---|---|---|---|---|---|---|---|---|---|---|---|
| | Age (years) | Sex* | Breed | Histology | BRAF$^{V595E}$ mutation | Treatment | IHC | Survival analysis | RNA-Seq | qPCR | Urine (ELISA) | Serum (ELISA) |
| PC1 | 11.2 | MC | Wire Fox Terrier | Prostate cancer | Mutation | Radical cystoprostatectomy | + | + | + | + | + | + |
| PC2 | 11.1 | MC | Yorkshire Terrier | Prostate cancer | Mutation | Radical cystoprostatectomy | + | + | + | + | + | + |
| PC3 | 12.7 | MC | English Cocker Spaniel | Prostate cancer | Mutation | Radical cystoprostatectomy | + | + | + | + | | |
| PC4 | 10.3 | MC | Border Collie | Prostate cancer | Mutation | Radical cystoprostatectomy | + | + | + | + | + | + |
| PC5 | 11.8 | MI | Border Collie | Prostate cancer | Wild-type | Radical cystoprostatectomy | + | + | + | + | | |
| PC6 | 11.9 | MC | Papillon | Prostate cancer | Mutation | Radical cystoprostatectomy | + | + | + | + | + | + |
| PC7 | 12.3 | MI | Miniature Dachshund | Prostate cancer | Mutation | Radical cystoprostatectomy | + | + | + | + | + | + |
| PC8 | 11.4 | MC | Labrador Retriever | Prostate cancer | Wild-type | Radical cystoprostatectomy | + | + | + | + | + | + |
| PC9 | 11.8 | MC | Maltese | Prostate cancer | Wild-type | Radical cystoprostatectomy | + | + | + | + | + | + |
| PC10 | 10.8 | MC | Miniature Dachshund | Prostate cancer | Mutation | Radical cystoprostatectomy | + | + | + | + | | |
| PC11 | 15.9 | MI | Miniature Dachshund | Prostate cancer | Wild-type | Radical cystoprostatectomy | + | + | | | | |
| PC12 | 9.3 | MC | Miniature Dachshund | Prostate cancer | Mutation | Radical cystoprostatectomy | + | + | | | | |
| PC13 | 11.9 | MC | Miniature Schnauzer | Prostate cancer | Mutation | Radical cystoprostatectomy | + | + | | | | |
| PC14 | 10.4 | MC | Pembroke Welsh Corgi | Prostate cancer | Mutation | Radical cystoprostatectomy | + | + | | | | |
| PC15 | 12.5 | MI | Miniature Schnauzer | Prostate cancer | Mutation | Radical cystoprostatectomy | + | + | | | | |
| PC16 | 11.7 | MC | Miniature Dachshund | Prostate cancer | Mutation | Radical cystoprostatectomy | + | + | + | + | + | + |
| PC17 | 9.0 | MC | Pug | Prostate cancer | Mutation | Radical cystoprostatectomy | + | + | + | + | + | + |
| PC18 | 11.0 | MC | Miniature Dachshund | Prostate cancer | Mutation | Radical cystoprostatectomy | + | + | + | + | + | + |
| PC19 | 9.9 | MC | Miniature Dachshund | Prostate cancer | Mutation | Piroxicam | | | + | + | + | + |
| PC20 | 11.9 | MC | Miniature Dachshund | Prostate cancer | Mutation | Firocoxib | | | | + | | |
| PC21 | 12.4 | MC | Miniature Dachshund | Prostate cancer | Mutation | Piroxicam | | | | + | + | + |
| PC22 | 11.4 | MC | Cavalier King Charles Spaniel | Prostate cancer | Mutation | Piroxicam | | | | + | + | + |
| PC23 | 8.2 | MC | Jack Russell Terrier | Prostate cancer | Mutation | Piroxicam | | | | + | + | + |
| PC24 | 12.6 | MC | Brussels Griffon | Prostate cancer | Mutation | Piroxicam | | | | | + | + |
| PC25 | 12.0 | MC | Papillon | Prostate cancer | Mutation | Piroxicam | | | | | + | + |
| PC26 | 13.3 | MC | Chihuahua | Prostate cancer | Wild-type | Piroxicam | | | | | + | + |
| PC27 | 8.9 | MI | Pembroke Welsh Corgi | Prostate cancer | Wild-type | Piroxicam | | | | | + | + |
| PC28 | 10.2 | MC | Miniature Dachshund | Prostate cancer | Mutation | Piroxicam | | | | | + | + |
| N1 | 8.1 | MI | Beagle | Normal | Wild-type | – | + | | + | + | + | + |
| N2 | 10.2 | MI | Beagle | Normal | Wild-type | – | + | | + | + | + | + |
| N3 | 9.0 | MI | Beagle | Normal | Wild-type | – | + | | + | + | + | + |
| N4 | 9.1 | MI | Beagle | Normal | Wild-type | – | + | | + | + | + | + |
| N5 | 4.5 | MI | Beagle | Normal | Wild-type | – | + | | | + | + | + |
| N6 | 4.5 | MI | Beagle | Normal | Wild-type | – | + | | | | + | + |
| N7 | 4.6 | MI | Beagle | Normal | Wild-type | – | + | | | | + | + |
| N8 | 4.6 | MI | Beagle | Normal | Wild-type | – | + | | | | + | + |
| N9 | 4.8 | MI | Beagle | Normal | Wild-type | – | + | | | | + | + |
| N10 | 8.8 | MI | Beagle | Normal | Wild-type | – | | | | | + | + |
| N11 | 10.4 | MC | Toy Poodle | Normal | Wild-type | – | | | | | + | |
| N12 | 8.9 | MC | Maltese | Normal | Wild-type | – | | | | | + | |
| N13 | 10.1 | MC | Miniature Dachshund | Normal | Wild-type | – | | | | | + | |
| N14 | 7.6 | MC | Papillon | Normal | Wild-type | – | | | | | + | |

*MC, male castrated; MI, male intact. [†]IHC, immunohistochemistry; RNA-Seq, RNA sequencing; qPCR, quantitative PCR; ELISA, enzyme-linked immunosorbent assay.

**Table S2. Characteristics of dogs with prostate cancer in the clinical trial.**

| Case ID | Age (years) | Sex* | Breed | TNM classification (at first visit) | $BRAF^{V595E}$ mutation | Treatment | Response† | Progression-free survival (days) | Overall survival (days) |
|---|---|---|---|---|---|---|---|---|---|
| M1 | 9.8 | MC | Papillon | T2N1M0 | Mutation | Mogamulizumab Piroxicam | PR | 105 | 296 |
| M2 | 11.5 | MC | Miniature Dachshund | T4N0M0 | Mutation | Mogamulizumab Piroxicam | SD | 66 | 381 |
| M3 | 7.2 | MC | Miniature Dachshund | T3N0M0 | Mutation | Mogamulizumab Piroxicam | SD | 505 | 1000 |
| M4 | 12.1 | MC | Miniature Dachshund | T4N0M0 | Mutation | Mogamulizumab Piroxicam | PR | 116 | 162 |
| M5 | 13 | MC | Miniature Dachshund | T3N0M0 | Wild-type | Mogamulizumab Piroxicam | SD | 287 | 368 |
| M6 | 9.6 | MI | Miniature Dachshund | T3N1M0 | Mutation | Mogamulizumab Piroxicam | PR | 209 | 232 |
| M7 | 12.9 | MC | Chihuahua | T4N1M0 | Wild-type | Mogamulizumab Piroxicam | SD | 57 | 147 |
| M8 | 14.8 | MC | Miniature Schnauzer | T4N0M0 | Wild-type | Mogamulizumab Piroxicam | SD | 208 | 458 |
| M9 | 12 | MC | Papillon | T4N1M1 | Wild-type | Mogamulizumab Piroxicam | SD | 42 | 86 |
| M10 | 11.6 | MC | Chihuahua | T3N1M1 | Mutation | Mogamulizumab Piroxicam | PR | 133 | 227 |
| M11 | 13.7 | MC | Toy Poodle | T4N1M0 | Wild-type | Mogamulizumab Piroxicam | PR | 63 | 404 |
| M12 | 12.6 | MC | Miniature Schnauzer | T2N0M0 | Mutation | Mogamulizumab Piroxicam | SD | 106 | 116 |
| M13 | 12.2 | MC | Miniature Dachshund | T3N0M0 | Mutation | Mogamulizumab Piroxicam | SD | 540 | 644 |
| M14 | 14.8 | MC | Miniature Dachshund | T2N0M0 | Mutation | Mogamulizumab Piroxicam | SD | 204 | > 812 |
| M15 | 13.4 | MC | Miniature Dachshund | T4N1M1 | Mutation | Mogamulizumab Piroxicam | SD | 238 | 345 |
| M16 | 12.4 | MI | Miniature Dachshund | T4N1M1 | Wild-type | Mogamulizumab Piroxicam | PD | 26 | 187 |
| M17 | 9.8 | MC | Miniature Dachshund | T4N1M0 | Mutation | Mogamulizumab Piroxicam | SD | 225 | 325 |
| M18 | 13.5 | MC | Toy Poodle | T4N0M0 | Mutation | Mogamulizumab Piroxicam | SD | 573 | > 667 |
| M19 | 13.6 | MC | Miniature Schnauzer | T4N1M1 | Mutation | Mogamulizumab Piroxicam | PR | 84 | 133 |
| M20 | 13.5 | MI | Pug | T4N1M0 | Wild-type | Mogamulizumab Piroxicam | PD | 21 | 136 |
| M21 | 13.8 | MC | Chihuahua | T2N0M0 | Mutation | Mogamulizumab Piroxicam | SD | 253 | 260 |
| M22 | 13.3 | MC | Pomeranian | T1N0M0 | Mutation | Mogamulizumab Piroxicam | PR | 377 | 970 |
| M23 | 13.2 | MC | Toy Poodle | T2N0M0 | Mutation | Mogamulizumam Piroxicam | SD | > 189 | > 189 |
| P1 | 8.5 | MC | Pembroke Welsh Corgi | T3N1M0 | Mutation | Piroxicam | PD | 22 | 22 |
| P2 | 11.1 | MI | French Bulldog | T4N0M0 | Mutation | Piroxicam | SD | 71 | 166 |
| P3 | 9.8 | MC | Miniature Dachshund | T4N1M1 | Mutation | Piroxicam | PD | 14 | 14 |
| P4 | 11.6 | MC | Miniature Dachshund | T3N1M0 | Wild-type | Piroxicam | SD | 46 | 68 |
| P5 | 11.8 | MC | Miniature Dachshund | T3N0M1 | Mutation | Piroxicam | SD | 65 | 65 |

| | | | | | | | | | |
|---|---|---|---|---|---|---|---|---|---|
| P6 | 12.1 | MC | Miniature Dachshund | T2N1M0 | Mutation | Piroxicam | PD | 30 | 206 |
| P7 | 11.4 | MC | Cavalier King Charles Spaniel | T4N1M1 | Wild-type | Piroxicam | PD | 6 | 6 |
| P8 | 11.4 | MC | Wire Fox Terrier | T4N0M0 | Mutation | Piroxicam | SD | 118 | 285 |
| P9 | 11.5 | MC | Yorkshire Terrier | T4N0M0 | Wild-type | Piroxicam | SD | 61 | 149 |
| P10 | 12.4 | MC | English Cocker Spaniel | T4N0M0 | Mutation | Piroxicam | SD | 94 | 148 |
| P11 | 12.1 | MI | Border Collie | T4N0M0 | Wild-type | Piroxicam | SD | 39 | 151 |
| P12 | 11.8 | MC | Papillon | T4N0M0 | Mutation | Piroxicam | SD | 57 | 92 |
| P13 | 13.8 | MC | Beagle | T2N0M0 | Wild-type | Piroxicam | PR | 182 | 263 |
| P14 | 12.3 | MC | Miniature Dachshund | T4N1M0 | Mutation | Piroxicam | PD | 29 | 29 |
| P15 | 12.5 | MC | Brussels Griffon | T4N1M0 | Mutation | Piroxicam | PD | 28 | 75 |
| P16 | 8.1 | MC | Jack Russel Terrier | T4N1M0 | Mutation | Piroxicam | PD | 16 | 103 |
| P17 | 8.6 | MC | Miniature Dachshund | T2N1M0 | Mutation | Piroxicam | PD | 26 | 56 |
| P18 | 12.2 | MC | Miniature Dachshund | T1N0M0 | Mutation | Piroxicam | SD | 182 | 229 |
| P19 | 12.4 | MI | Miniature Schnauzer | T3N1M1 | Mutation | Piroxicam | SD | 42 | 77 |
| P20 | 15.7 | MC | Miniature Dachshund | T2N0M0 | Mutation | Piroxicam | SD | 210 | 389 |
| P21 | 8.9 | MI | Pembroke Welsh Corgi | T4N1M1 | Wild-type | Piroxicam | SD | 85 | 99 |
| P22 | 13 | MC | Toy Poodle | T4N0M0 | Mutation | Piroxicam | PR | 139 | 468 |
| P23 | 12.2 | MC | Miniature Dachshund | T2N0M0 | Mutation | Piroxicam | SD | 80 | 86 |

*MC, male castrated; MI, male intact. †SD, stable disease; PR, partial response; PD, progressive disease.

.

**Table S3. Characteristics of human patients with prostate cancer.**

| Case ID | Age (years) | Sex* | Pathological diagnosis | Gleason score | TNM classification | Minimum stage grouping | Foxp3$^{+}$ cells (cells/HPF)† | CCR4$^{+}$ cells (cells/HPF)† |
|---|---|---|---|---|---|---|---|---|
| hPC1 | 74 | M | Adenocarcinoma of prostate | 3+4=7/10 | pT2apN0pMX | II | 2.2 | 5.2 |
| hPC2 | 60 | M | Adenocarcinoma of prostate | 3+4=7/10 | pT2cpNXpMX | II | 8 | 16.4 |
| hPC3 | 52 | M | Adenocarcinoma of prostate | 3+4=7/10 | pT2bpN0pMX | II | 5 | 0.8 |
| hPC4 | 59 | M | Adenocarcinoma of prostate | 3+4=7/10 | pT2cpNXpMX | II | 17 | 29.6 |
| hPC5 | 61 | M | Adenocarcinoma of prostate | 3+4=7/10 | pT3bpN0pMX | III | 5.2 | 4.2 |
| hPC6 | 75 | M | Adenocarcinoma of prostate | 3+4=7/10 | pT2cpN0pMX | II | 14.6 | 51.6 |
| hPC7 | 54 | M | Adenocarcinoma of prostate | 3+4=7/10 | pT2apN0pMX | II | 4.4 | 18.4 |
| hPC8 | 57 | M | Adenocarcinoma of prostate | 3+4=7/10 | pT2bpN0pMX | II | 19.8 | 46.6 |
| hPC9 | 62 | M | Adenocarcinoma of prostate | 3+4=7/10 | pT3bpNXpMX | III | 4.8 | 2.2 |
| hPC10 | 61 | M | Adenocarcinoma of prostate | 3+4=7/10 | pT3bpNXpMX | III | 12.4 | 9.8 |
| hPC11 | 58 | M | Adenocarcinoma of prostate | 3+4=7/10 | pT2cpN0pMX | II | 12.6 | 10.2 |
| hN1 | 62 | M | Normal | – | – | – | 3 | 0.8 |
| hN2 | 54 | M | Normal | – | – | – | 1.2 | 0.2 |
| hN3 | 65 | M | Normal | – | – | – | 1 | 4.8 |
| hN4 | 66 | M | Normal | – | – | – | 1.4 | 1.4 |
| hN5 | 70 | M | Normal | – | – | – | 2.2 | 2 |
| hN6 | 65 | M | Normal | – | – | – | 2 | 2.6 |

*M, male. †HPF, high power field.

**Table S4. Primer pair sequences used for quantitative RT-PCR.**

| Primer set | GenBank accession number | | Primer sequence (5′–3′) |
|---|---|---|---|
| IL-10 | NM_001003077 | Forward | CGA CCC AGA CAT CAA GAA CC |
| | | Reverse | CAC AGG GAA GAA ATC GGT GA |
| CCL3 | NM_001005251 | Forward | CAA GCA GAT TCC ACG CAA GGT |
| | | Reverse | TAA TAC CGG GCT TGG AGC AT |
| CCL4 | NM_001005250 | Forward | CGT CCT TTC TCT CCT TGT GC |
| | | Reverse | GAA TCT TCC GCA GGG TGT AA |
| CCL5 | NM_001003010 | Forward | GGT CTC CGC AGC TAC CTT T |
| | | Reverse | AAA GCA GCA GGG TGT GGT |
| CCL7 | NM_001010960 | Forward | CCC ATC CAG AAG CTG AAG AG |
| | | Reverse | CGT CCT TAG CCA GTT TGG TC |
| CCL8 | NM_001005255 | Forward | GTC CTT GCT CAG CCA GAT TC |
| | | Reverse | ACT GGC TGT TGG TGA TCC TC |
| CCL13 | NM_001003966 | Forward | GCC CTA TTC ACT TGC TGC TT |
| | | Reverse | AAT CCT GGA CCC ATT TCT CC |
| CCL14 | XM_537723 | Forward | TCA CGA GGA CCT TAC CAT CC |
| | | Reverse | GGC CAT TTT TGG TGA TGA AG |
| CCL17 | NM_001003051 | Forward | GGC TGA CAA GGT GGT ACA AGA CTT C |
| | | Reverse | CAG ATG GAC TTG CCT TGG ACA G |
| CCL22 | XM_003433778 | Forward | TAT GGT GCC AAC GTG GAA GA |
| | | Reverse | GAT CTC CCG ATC CTT GAC AGT TAG |
| CCL24 | NM_001003967 | Forward | CCT GCT GCA TGT TCT TCA TTT C |
| | | Reverse | TTC TGG TTC TTC TTG GTG GTG A |
| CCL28 | NM_001005257 | Forward | CAG ACA GGA CTC ACT CTC GCT CTC |
| | | Reverse | TGT GAA ACC TCA GTG CAA CAG CTA |
| CXCL8 | NM_001003200 | Forward | CTT CCA AGC TGG CTG TTG CTC |
| | | Reverse | TGG GCC ACT GTC AAT CAC TCT C |
| CXCL10 | NM_001010949 | Forward | ATT GAG ATG ATT CCT GCA AGT |
| | | Reverse | TCA GAC ATC TTT TCT CCC CAC TC |
| CXCL13 | XM_845089 | Forward | GGG TGC CCA AAA AGA GAA ATC |
| | | Reverse | GAT GGG AGG GTT CAA GCA TAC A |
| CXCL16 | XM_014113226 | Forward | GAG AGC CAG AAG CAG CAG AT |
| | | Reverse | GTG ACT GCT CCC TCC TCT TG |
| CX3CL1 | AB648939 | Forward | CTT CCT TGG CCT CCT CTT CT |
| | | Reverse | GGC ACC AGG ACA TAC GAG TT |
| RPL13A | AJ388525 | Forward | GCC GGA AGG TTG TAG TCG T |
| | | Reverse | GGA GGA AGG CCA GGT AAT TC |
| RPL32 | XM_848016 | Forward | TGG TTA CAG GAG CAA CAA GAA A |
| | | Reverse | GCA CAT CAG CAG CAC TTC A |

IL, interleukin; RPL, ribosomal protein L.
